# Effect of percutaneous ventricular assisted device on post-cardiac arrest myocardial dysfunction in swine model with prolonged cardiac arrest

**DOI:** 10.1101/2023.08.01.551568

**Authors:** Takahiro Nakashima, Mohamad Hakam Tiba, Cindy H. Hsu, Adam L. Gottula, Brendan M. McCracken, Nicholas L. Greer, Traci A. Cramer, Nadia R. Sutton, Kevin R. Ward, Robert W Neumar

## Abstract

**Background:** Percutaneous left ventricular assist device (pLVAD) can provide hemodynamic support during and after cardiac arrest, but it remains unclear if pLVAD reduces post-cardiac arrest myocardial dysfunction.

**Methods:** This is an analysis of a subset of animals that achieved return of spontaneous circulation (ROSC) in a study comparing pLVAD, transient aortic occlusion (AO), or both during cardiopulmonary resuscitation (CPR) after prolonged cardiac arrest. pLVAD, AO, or both were initiated after 24 minutes of ventricular fibrillation cardiac arrest (8 min no-flow and 16 min mechanical CPR). AO was discontinued post-ROSC, and pLVAD support or standard care were continued. Beginning 60 minutes post-ROSC, pLVAD support was weaned to <1.0 L/min while maintaining a mean arterial pressure >70 mmHg, and subsequently removed at 240 minutes when feasible. The primary outcome was the recovery of cardiac index (CI), stroke volume index (SVI), and left ventricular ejection fraction (LVEF) at 240 minutes post-ROSC. Data are shown as mean (standard error).

**Results:** Seventeen animals achieved ROSC without complication and were included in this analysis (pLVAD group, n = 11 and standard care group, n = 6). For the primary outcome, the pLVAD group had significantly higher CI of 4.2(0.3) vs. 3.1(0.4) L/min/m2 (p=0.043) and LVEF 60(3) vs. 49(4) % (p=0.029) at 240 minutes after ROSC, respectively, when compared with the standard care group, while SVI was not significant difference (2[3] vs. 23[4] mL/min/m^2^, *p*=0.054). During the first 60 minutes after ROSC with maximum pLVAD flow, the pLVAD group had significantly higher coronary perfusion pressure (62[4] vs. 47[5] mmHg, p=0.019), lower LV stroke work index (3.9[3.0] vs. 14.9[4.4] cJ/m2, p=0.043), and lower total pulmonary resistance index (13.2[4.8] vs. 21.5[14.4] Wood Unit, p=0.001).

**Conclusion:** These results suggest that early pLVAD support after ROSC is associated with better recovery myocardial function compared to standard care after prolonged cardiac arrest.

## Introduction

The survival rate for patients with prolonged cardiac arrest remains extremely low, even with recent advancements in post-cardiac arrest care.^1–3^ 63-72% of patients that achieve return of spontaneous circulation (ROSC) after out-of-hospital cardiac arrest (OHCA) die before hospital discharge.^4, 5^ This high post-resuscitation mortality primarily results from brain injury, but death within the first 24 hours after ROSC is typically attributed to refractory shock and multiple organ failure due to myocardial dysfunction.^4, 6–8^ Post-cardiac arrest myocardial dysfunction is reported to occur in 49-68% of patients after ROSC.^7–10^

A micro-axial percutaneous left ventricular assist device (pLVAD), which actively pumps blood flow from the LV to the ascending aorta, has become widely used in patients with cardiogenic shock. This device has been reported to reduce the LV load, as well as increase coronary perfusion in animal experiments.^11, 12^ Several studies have reported a potential role for pLVAD in patients with cardiac arrest.^13–15^ Recently, our team demonstrated that adding aortic balloon occlusion to pLVAD had significantly higher coronary and cerebral perfusion pressure during cardiopulmonary resuscitation (CPR) in a swine model with prolonged cardiac arrest.^16^ In addition, an animal study, using dogs with a myocardial infarction model, reported that LV mechanical unloading by pLVAD in the acute phase of MI reduced infarct size. However, there is a critical knowledge gap of evidence on the impact of pLVAD on post-cardiac arrest myocardial dysfunction. We hypothesized that the use of pLVAD would promote the recovery of myocardial function even after prolonged cardiac arrest.

To address this hypothesis, we examined the effectiveness of early pLVAD support on the recovery of myocardial function after prolonged cardiac arrest using the swine model of prolonged cardiac arrest. In addition, we assessed the trajectory of cardiac function and hemodynamics recovery post-ROSC.

## Methods

This study was the prespecified analysis of post-ROSC hemodynamics data from the RCT comparing three types of CPR methods. Twenty-four Yorkshire mix swine weighing 45 – 55kg (males: n=12) were subjected to cardiac arrest and underwent CPR and randomly assigned to one of three CPR groups; swine that received pLVAD with (n=8) and without (n=8) transient aortic balloon occlusion during CPR or transient aortic balloon occlusion alone during CPR (n=8). If ROSC was achieved, the aortic balloon was immediately deflated, and the two arms of post-cardiac arrest care were continued for 240 minutes; 1) pLVAD support group or 2) Standard care group (control).

This study was conducted in compliance with the Guide for Care and Use of Laboratory Animals (US National Institutes of Health Publication No. 85-23, National Academy Press, Washington D.C revised in 2011. All methods and procedures were approved by the University of Michigan Institutional Animal Care and Use Committee and followed the Animals in Research: Reporting In Vivo Experiments (ARRIVE) guidelines.^17^

### Surgical preparation

Anesthesia was induced using a mixture of intramuscular Telazol/xylazine. Animals were intubated endotracheally and mechanically ventilated with 7.5-10 mL Kg^−1^ tidal volume. Anesthesia was maintained by inhalant isoflurane (1-2%) delivered in 0.3-0.4 oxygen with balanced nitrogen. Respiratory rate was adjusted to titrate an end-tidal CO_2_ (PetCO_2_) to 35– 45 mmHg (Biopac Data Acquisition System. Biopac Inc. Goleta, CA). Core body temperature was maintained using a blanket to a target of 37 – 38 °C.

Both common femoral arteries were cannulated using a 9 Fr introducer (Arrow International Inc., Reading, PA) to place a pLVAD or the aortic balloon occlusion catheter. The right common carotid artery was cannulated with a catheter to the aortic arch to monitor mean arterial pressure (MAP). An ultrasonic flow probe (3PS. Transonic, Ithaca, NY) was placed around the left common carotid artery to measure blood flow (CCA-Flow). A triple lumen central venous catheter (Arrow International Inc., Reading, PA) was placed in the right internal jugular vein to the right atrium for right atrial pressure measurement, fluid infusion, and blood sampling. The right external jugular vein was cannulated with a 9Fr introducer via ultrasound-guided Seldinger technique for the placement of an 8 Fr pulmonary artery (PA) catheter (Edwards LifeSciences LLC, Irvine, CA) to continuously monitor cardiac output and mixed venous oxygen saturation (SVO_2_). A multichannel Data Acquisition System (MP150, Biopac, Goleta, CA) recorded all hemodynamic monitoring throughout the cardiac arrest period.

### Prolonged cardiac arrest and cardiopulmonary resuscitation protocol

Baseline data and blood samples were collected 15 min following instrumentation. Prior to cardiac arrest, pLVAD or aortic balloon occlusion catheter was advanced through the common femoral artery according to assigned CPR groups. A 14 Fr pLVAD, Impella-CP (Abiomed, Danvers, MA), was placed in the LV under the fluoroscopy guidance and positioned to ensure a pLVAD flow rate of P7 or greater. A 7F aortic occlusion catheter, Fogarty (Edwards Lifesciences, Irvine, CA), was placed at a pre-measured distance to the diaphragm (approximately 50-55 cm). Balloon inflation or deflation was verified by femoral arterial pressure pulsatility on the same side.

Ventricular fibrillation (VF) was induced with the pacing wire using a 9 V battery. VF was verified by ECG, and loss of pulsatile AoBP and noted as time zero (T0). Anethsethia and ventilation were paused. VF was left untreated for 8 minutes (T8). CPR was started at T8 with a mechanical chest compression device (LUCAS-II, Stryker Medical, Portage MI) which provided 100 compressions per minute at a depth of 5 cm, and ventilation was resumed with a respiratory rate of 10 /minute at the previous tidal volume and 100% FiO_2_. Epinephrine (0.015mg/kg) was administered at T14 and T18. Sixteen minutes after mechanical CPR initiation (T24), a pre-implemented pLVAD or aortic occlusion catheter was activated under continuous mechanical CPR. pLVAD flow was increased to the maximum possible rate without a suction alarm. Four minutes after devices were initiated (T28), the first 200 J biphasic defibrillation (Medtronic LIFEPAK 20e, Minneapolis MN) was attempted. If ROSC, defined as an organized electrical rhythm or pulse with measured arterial pressure for 2 minutes, was achieved, a post-cardiac arrest care phase was initiated. Otherwise, CPR and interventions were continued for 8 more minutes with a single defibrillation attempt every subsequent 4 minutes. If ROSC was not achieved after any defibrillation attempt (T36 of total arrest time), the experiment was terminated. We reported the CPR quality elsewhere.

### Post-cardiac arrest care protocol

Animals that achieved ROSC were immediately started on vasopressors and isoflurane anesthesia. The aortic occlusion catheter was deflated and removed 2 minutes after ROSC was confirmed regardless of hemodynamic stability. Animals were continued on pLVAD supportive care or standard of care with vasopressors for 240 minutes according to treatment arms. The flow rate of pLVAD was further increased up to the maximum possible flow rate immediately after ROSC and continued for an additional 60 minutes before gradual weaning was attempted. We confirmed the proper placement of pLVAD using fluoroscopy and adjusted the position. The pulmonary artery catheter was reimplemented after ROSC was sustained for more than 20 minutes. Epinephrine (0-1.0 mcg/kg/min) and norepinephrine (0-1.0 mcg/kg/min) were used for pressor support as a target of MAP ≥70 mmHg, total cardiac output ≥2.0 L/min, and central venous oxygen saturation (SVO_2_) >55%. If a stable MAP ≥70mmHg and SVO_2_ >55% could be maintained, the vasopressors were actively reduced per 0.02 −0.05 mcg/kg/min. Epinephrine was decreased prior to norepinephrine if a stable total cardiac output ≥2.0 L/min could be maintained. If MAP ≥70 mmHg and SVO_2_ >55% are maintained, but epinephrine could not be reduced due to low total cardiac output, norepinephrine was reduced. In animals with pLVAD supportive care, pLVAD flow was maintained at maximum flow rate until 60 minutes after ROSC. The weaning ability from Impella was assessed 60 minutes after ROSC with a progressively decreased Impella flow rate. Weaning was deemed successful if a stable MAP > 70 mmHg could be maintained with negligible Impella flow (flow rate of P1 with <1.0 L/min) or off. After completed the study, animals were euthanized under anesthesia.

### Data and sample collection

All transthoracic echocardiography (TTE) studies were performed by the same board-certificated cardiologist (T.N) using a commercially available ultrasound machine with 2.18 Hz transducer (M9, Mindray, Mahwah, NJ) at baseline, at ROSC, and every 30 minutes after ROSC. In brief, at least three consecutive cardiac cycles were stored and averaged for all measurements. We evaluated LV end-systolic volume (LVESV), LV end-diastolic volume (LVEDV), LVEF, and velocity time integral of the LV outflow tract (LVOT-VTI). We measured the anterior-posterior (short axis and long axis) dimension and calculated the left ventricular volume (LVV) by the Teichholz formula: LVV = 7D^3^ / (2.4+D).

Hemodynamic data were continuously monitored by PA catheter, including coronary perfusion pressure (CPP), cardiac output / cardiac index (CI) by thermodilution technique, stroke volume (SV), stroke volume index (SVI), pulmonary artery wedge pressure (PCWP), pulmonary artery pressure (PAP) and central venous pressure (CVP). We also calculated body surface (BSA),^18^ systemic vascular resistance index (SVRI), total pulmonary resistance index (TPRI), PA compliance, and LV stroke work index (LVSWI) as follows;

CPP (mmHg^2^) = diastolic arterial pressure – PCWP

BSA(m^2^) = 0.0734 × body weight (kg)^0.656^

SVRI (Wood Unit) = (MAP – CVP)/CO /BSA,

PVRI (Wood Unit) = (mean PAP – PCWP) /CI /BSA,

TPRI (Wood Unit) = mean PAP /CO /BSA,

PA compliance = SV/pulmonary artery pulse pressure,

LVSWI (cJ/m^2^) = (MAP – PCWP) × native SV × 0.0133322 /BSA.

Native cardiac output was calculated using the difference between total cardiac output calculated by pulmonary artery catheter and pLVAD flow calculated by the Impella controller.

### Outcome measure

The primary outcome was the recovery of CI and SVI measured by PA catheter and LVEF measured by TTE at 240 minutes after ROSC.

### Statistical analysis

We compared the recovery of cardiac function 240 minutes after ROSC as a primary outcome between the group treated with pLVAD and the group treated with standard care. Then, to examined how early pLVAD support affect heart, we compared hemodynamics between the groups focusing on the early period within 60 minutes after ROSC when maximum pLVAD flow was maintained. For baseline characteristics, we compared distributed variables using a t-test, and the results are presented as the mean (standard deviation [SD]). Categorical variables were presented as count (percentage), and compared using the χ² test. The longitudinal data was presented as the mean (standard error [SE]), and analyzed by using a mixed effect model for repeated measures, which applied an autoregressive covariance structure to account for the correlation between subjects. Group effect at a given time point was estimated and tested in models with adjustment for multiple comparisons using the Bonferroni method. An α value of less than 0.05 was considered to indicate a significant difference. Statistical analyses were performed with R statistical software V.4.0.2 (R Foundation for Statistical Computing, Vienna, Austria).

## Results

Among a total of 24 swine, 20 swine (83%) achieved ROSC after prolonged cardiac arrest. Of these, two animals were excluded for severe complications by mechanical CPR [massive pneumothorax (n=1) and pericardial effusion (n=1)]. One animal with standard of care was hemodynamically unstable with MAP below 30 mmHg despite a maximum dose of inotrope and died within 30 minutes after ROSC. Finally, 17 swine were included in the analysis of cardiac function after ROSC; pLVAD group, n=11 and Standard care group, n=7 (**Figure 1**). Table 1 lists baseline physiologic variables for both groups. There was no difference between the groups at baseline.

**Figure 1.**
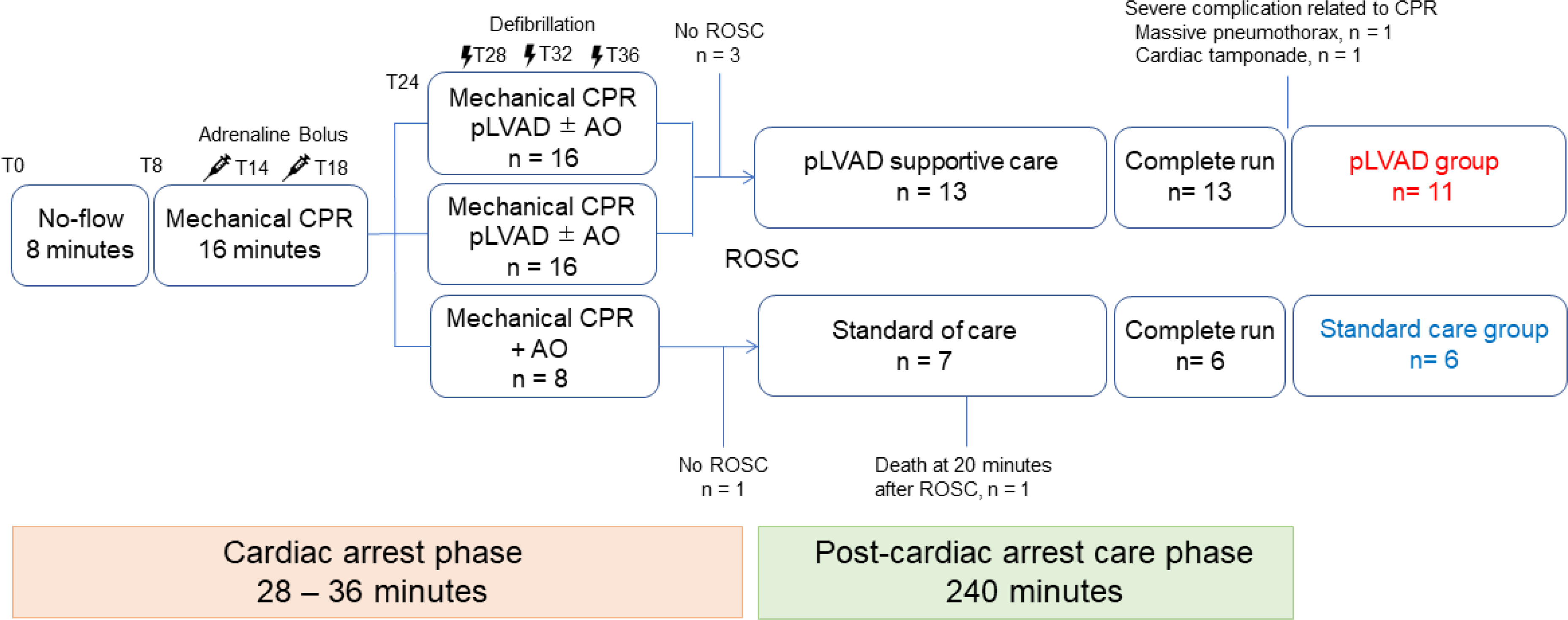
Study design. AO denotes aortic occlusion; CA, cardiac arrest; CPR cardiopulmonary resuscitation; pLVAD, percutaneous left ventricular assist device; and ROSC, return of spontaneous circulation.

**Table 1.**
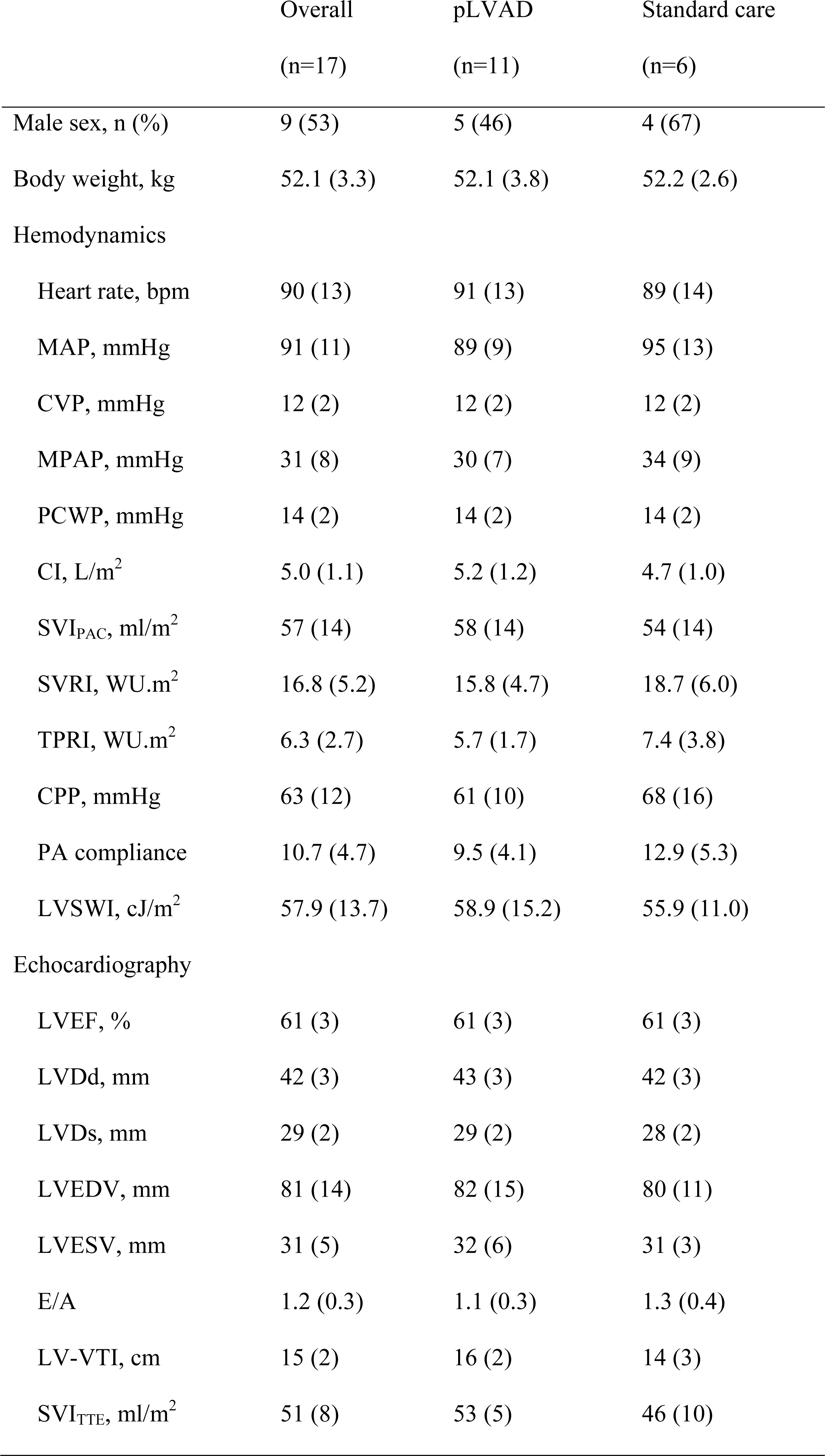
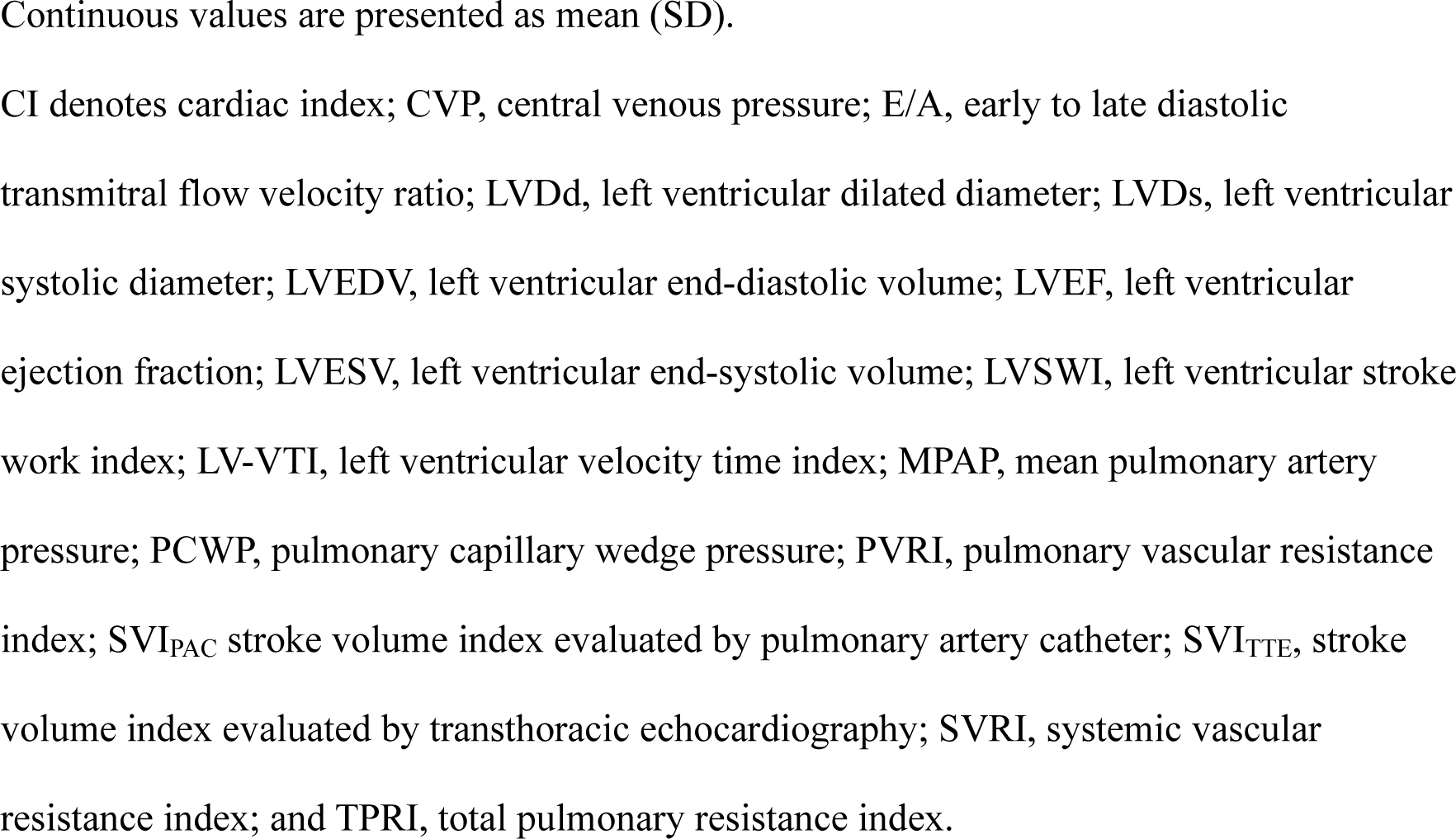
Baseline characteristics.

|  | Overall<br>(n=17) | pLVAD<br>(n=11) | Standard care<br>(n=6) |
| --- | --- | --- | --- |
| Male sex, n (%) | 9 (53) | 5 (46) | 4 (67) |
| Body weight, kg | 52.1 (3.3) | 52.1 (3.8) | 52.2 (2.6) |
| Hemodynamics |  |  |  |
| Heart rate, bpm | 90 (13) | 91 (13) | 89 (14) |
| MAP, mmHg | 91 (11) | 89 (9) | 95 (13) |
| CVP, mmHg | 12 (2) | 12 (2) | 12 (2) |
| MPAP, mmHg | 31 (8) | 30 (7) | 34 (9) |
| PCWP, mmHg | 14 (2) | 14 (2) | 14 (2) |
| CI, L/m<sup>2</sup> | 5.0 (1.1) | 5.2 (1.2) | 4.7 (1.0) |
| SVI<sub>PAC</sub>, ml/m<sup>2</sup> | 57 (14) | 58 (14) | 54 (14) |
| SVRI, WU.m<sup>2</sup> | 16.8 (5.2) | 15.8 (4.7) | 18.7 (6.0) |
| TPRI, WU.m<sup>2</sup> | 6.3 (2.7) | 5.7 (1.7) | 7.4 (3.8) |
| CPP, mmHg | 63 (12) | 61 (10) | 68 (16) |
| PA compliance | 10.7 (4.7) | 9.5 (4.1) | 12.9 (5.3) |
| LVSWI, cJ/m<sup>2</sup> | 57.9 (13.7) | 58.9 (15.2) | 55.9 (11.0) |
| Echocardiography |  |  |  |
| LVEF, % | 61 (3) | 61 (3) | 61 (3) |
| LVDd, mm | 42 (3) | 43 (3) | 42 (3) |
| LVDs, mm | 29 (2) | 29 (2) | 28 (2) |
| LVEDV, mm | 81 (14) | 82 (15) | 80 (11) |
| LVESV, mm | 31 (5) | 32 (6) | 31 (3) |
| E/A | 1.2 (0.3) | 1.1 (0.3) | 1.3 (0.4) |
| LV-VTI, cm | 15 (2) | 16 (2) | 14 (3) |
| SVI<sub>TTE</sub>, ml/m<sup>2</sup> | 51 (8) | 53 (5) | 46 (10) |

Continuous values are presented as mean (SD).
CI denotes cardiac index; CVP, central venous pressure; E/A, early to late diastolic transmitral flow velocity ratio; LVDd, left ventricular dilated diameter; LVDs, left ventricular systolic diameter; LVEDV, left ventricular end-diastolic volume; LVEF, left ventricular ejection fraction; LVESV, left ventricular end-systolic volume; LVSWI, left ventricular stroke work index; LV-VTI, left ventricular velocity time index; MPAP, mean pulmonary artery pressure; PCWP, pulmonary capillary wedge pressure; PVRI, pulmonary vascular resistance index; $SVI_{PAC}$ stroke volume index evaluated by pulmonary artery catheter; $SVI_{TTE}$ , stroke volume index evaluated by transthoracic echocardiography; SVRI, systemic vascular resistance index; and TPRI, total pulmonary resistance index.

### Echocardiographic variables at return of spontaneous circulation

The results of TTE variables at ROSC are shown in **Figure 2**. All variables were markedly reduced in both groups. LVEF, LVEDV, and LVESV were higher in the pLVAD group than in the Standard of care group, but there were no statistically significant differences with the exception of native SVI estimated by TTE (6 [10] vs. 20 [5] mL/m^2^, *p* = 0.007).

**Figure 2.**
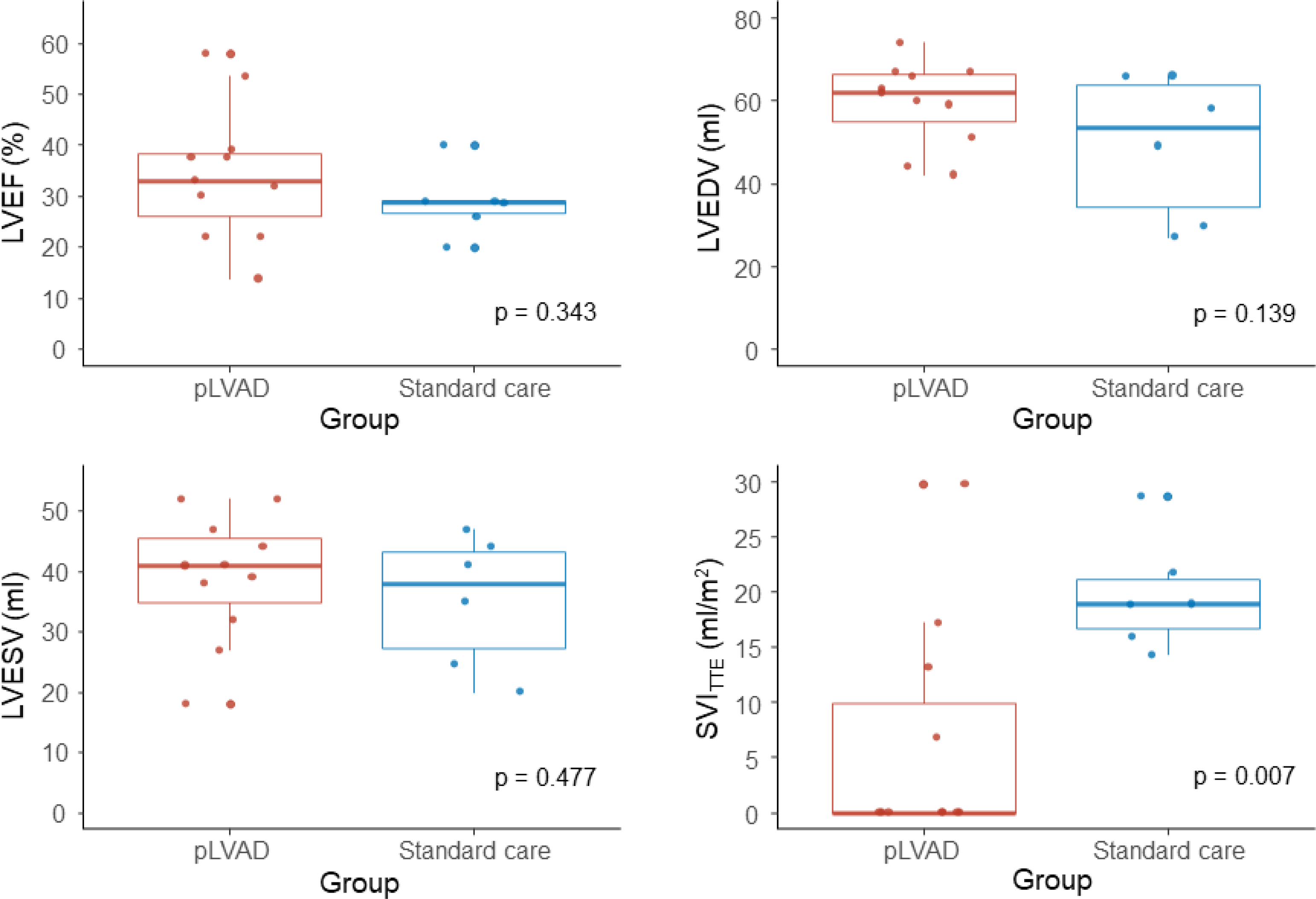
Transthoracic echocardiography variables at return of spontaneous circulation. LVEDV denotes left ventricular end-diastolic volume; LVEF, left ventricular ejection fraction; LVESV, left ventricular endo-systolic volume; pLVAD, percutaneous left ventricular assist device; and SVI_TTE_, stroke volume index evaluated by transthoracic echocardiography.

### Recovery of cardiac function after ROSC

The maximum pLVAD flow rate was 2.6 (0.1) L/min within 60 minutes after ROSC, which was 84 (26) % of total cardiac output at 30 minutes and 73 (24) % at 60 minutes after ROSC (**Figure 3**). At 240 minutes after ROSC, eight animals were removed from pLVAD and remaining three animals were minimally supported with flow rate of P1 with 0.2 (0.3) L/min. As depicted in **Figure 4**, there was a marked decrease in CI, SVI, and LVEF in the pLVAD groups and Standard care group at 30 minutes post ROSC, with mean values of 3.0 (0.3) vs. 2.3 (0.4) L/min/m^2^, 19 (3) vs. 15 (4) mL/min/m^2^ and 40 (3) vs. 32 (4) %, respectively. These values gradually recovered throughout the study, ultimately reaching 84 (20) %, 57 (19) %, and 99 (11) % of baseline in the pLVAD group and 64 (6) %, 45 (18) %, and 64 (6) % of baseline in the Standard care group, respectively. For primary outcomes, CI as evaluated by the PA catheter (4.2 [0.3] vs. 3.1 [0.4] L/min/m^2^, *p*=0.043) and LVEF (60 [3] vs. 49 [4] %, *p*=0.029) were significantly higher in the pLVAD group at 240 minutes after ROSC compared with the Standard care group (**Table 2**). SVI was also higher in the pLVAD group than in the Standard care group, however, without statistically significant difference (32 [3] vs. 23 [4] mL/min/m^2^, *p*=0.054).

**Figure 3.**
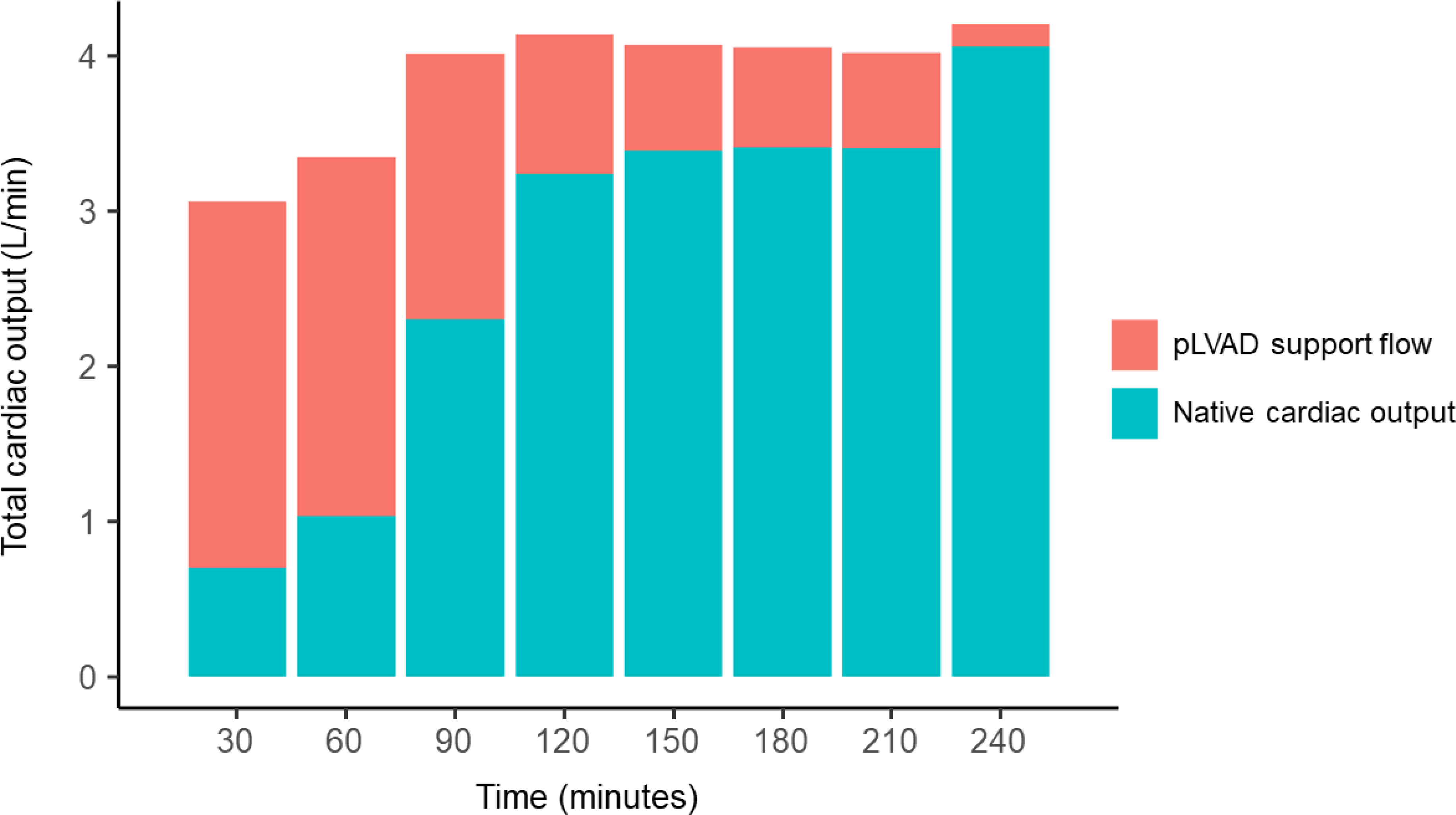
pLVAD support flow and native cardiac output. pLVAD denotes percutaneous left ventricular assist device.

**Figure 4.**
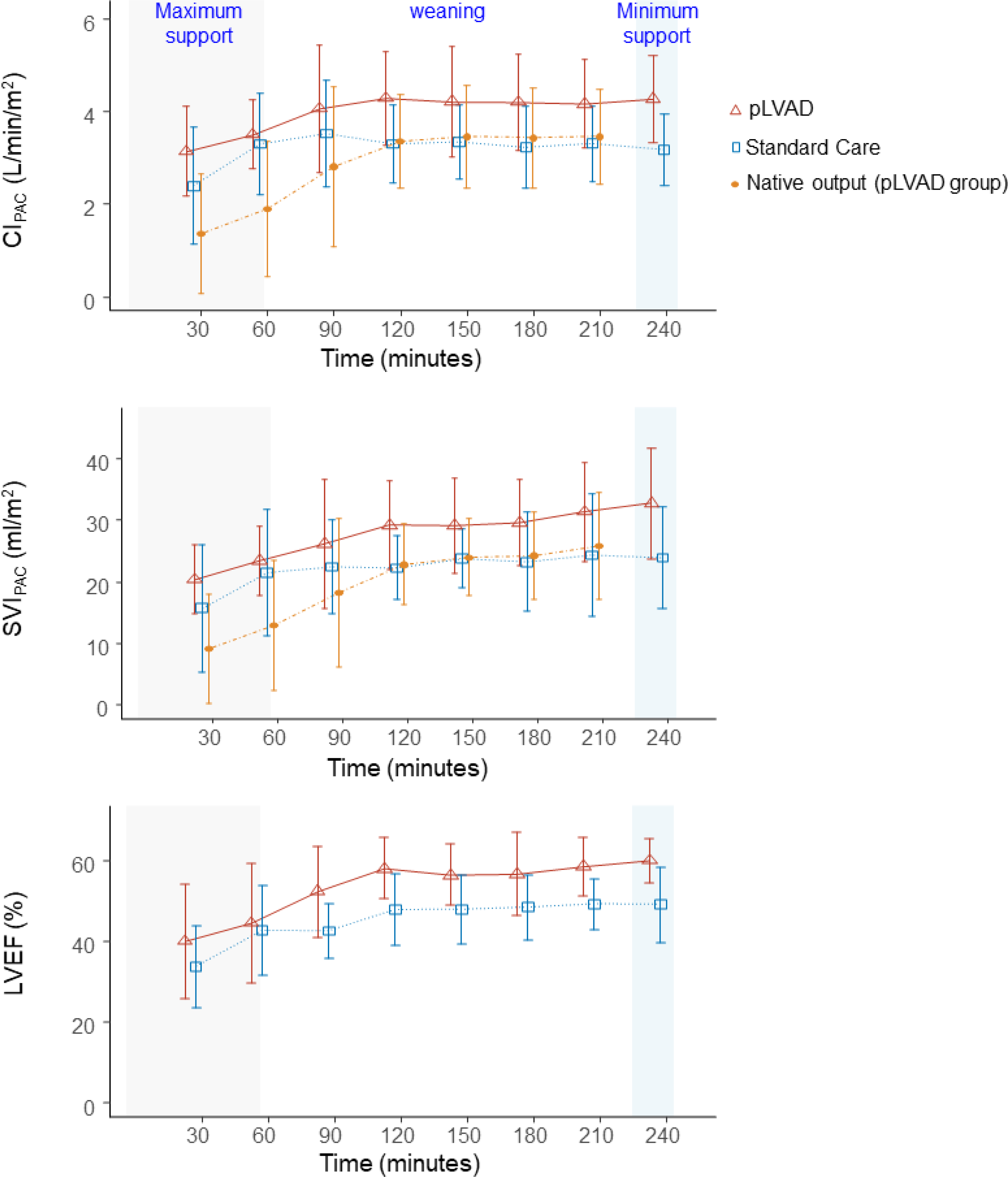
Longitudinal analyses of cardiac output measurements. Red and blue lines indicate pLVAD group and Standard care group, respectively. CI denotes cardiac index; LVEF, left ventricular ejection fraction; pLVAD, percutaneous left ventricular assist device; and SVI_PAC_ stroke volume index evaluated by pulmonary artery catheter.

**Table 2.** Comparison of cardiac output measurements at 240 minutes after resuscitation.

|  | pLVAD<br>(n=11) | Standard care<br>(n=6) | Mean absolute<br>Difference | p-value |
| --- | --- | --- | --- | --- |
| CI, L/m<sup>2</sup> | 4.2(0.3) | 3.1(0.4) | 1.1(0.5) | 0.043 |
| SVI<sub>PAC</sub>, mL/m<sup>2</sup> | 32(3) | 23(4) | 9(5) | 0.054 |
| LVEF, % | 60(3) | 49(4) | 11(5) | 0.029 |

Continuous values are presented as mean (SE).
CI denotes cardiac index; LVEF, left ventricular ejection fraction; SE, standard error; and SVI<sub>PAC</sub> stroke volume index evaluated by pulmonary artery catheter.

### Longitudinal analysis of hemodynamics

**Figures 5** show the trend of hemodynamics after ROSC. MAP was constantly maintained at greater than 70 mmHg in both groups throughout the study. There were no statistically significant differences in pressure variables and HR between the treatment groups at any time.

**Figure 5.**
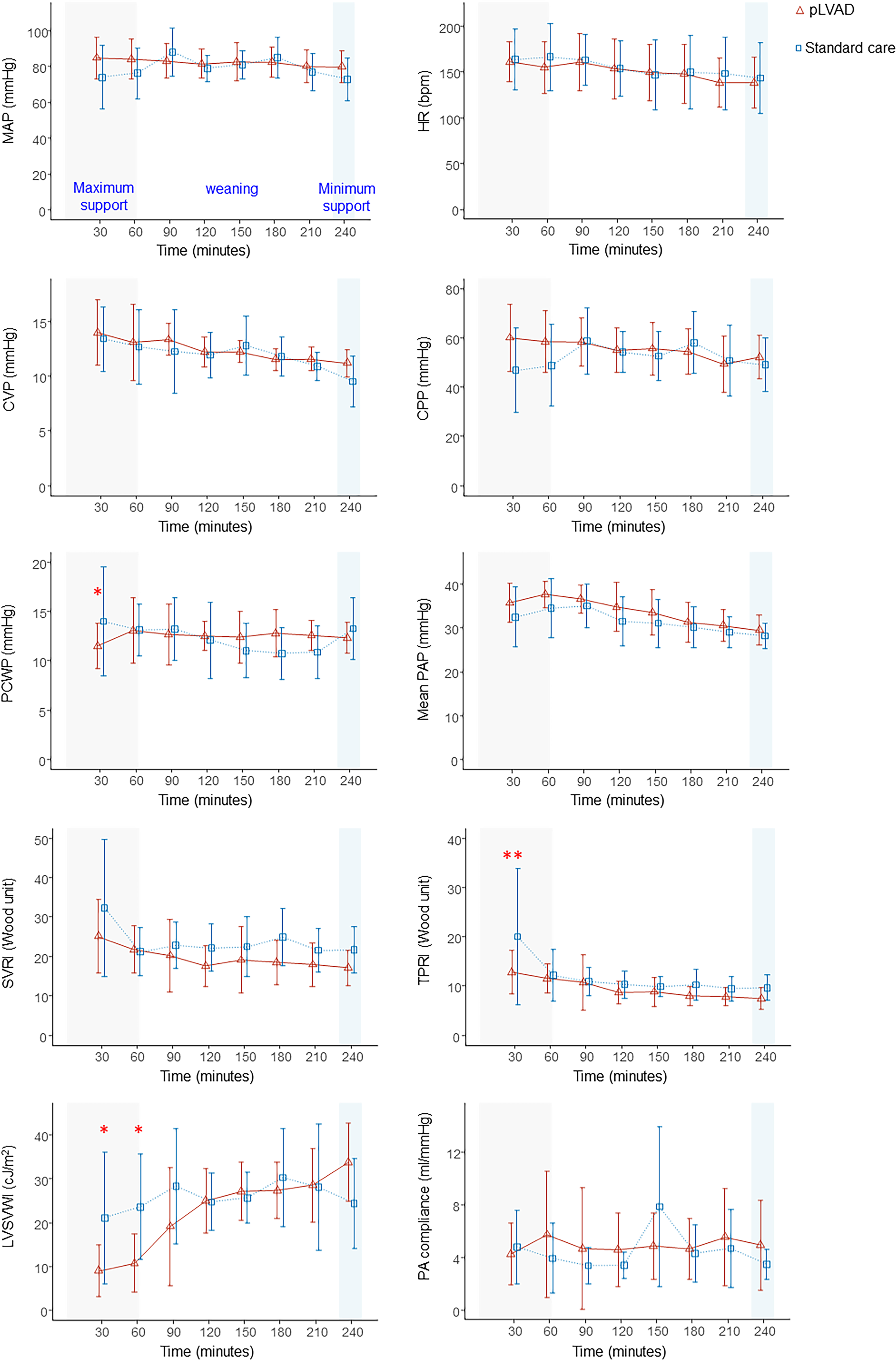
Longitudinal analyses of vascular resistance and stroke work after resuscitation. Red and blue lines indicate pLVAD group and Standard care group, respectively. * statistically significance (p <0.05) ** statistically significance (p<0.01) CPPc denotes coronary perfusion pressure; CVP, central venous pressure; HR, heartrate; LVSWI denotes left ventricular stroke work index; MAP, mean arterial pressure; PA, pulmonary artery; PAP pulmonary artery pressure; PCWP, pulmonary capillary wedge pressure; pLVAD, percutaneous left ventricular assist device; SVRI, systemic vascular resistance; and TPRI, total pulmonary resistance index.

Focusing on the early period within 60 minutes after ROSC when maximum pLVAD flow was maintained, the pLVAD group had significantly higher CPP than the Standard care group at 30 minutes (62 [4] vs. 47 [5] mmHg, *p*=0.019). The pLVAD group had significantly lower PCWP at 30 minutes (11 [1] vs. 14 [1] mmHg, *p*=0.041), and LVSWI at 30 minutes (3.9 [3.0] vs. 14.9 [4.4] cJ/m^2^, *p*=0.043) and 60 minutes (4.9 [3.3] vs. 18.2 [4.1] cJ/m^2^, *p*=0.013). Also, the pLVAD group had a significantly lower TPRI at 30 minutes with mean values of 13.2 (4.8) vs. 21.5 (14.4) Wood Unit (*p*=0.001). There was no significant difference in SVRI and PA compliance between the two groups. For vasopressor, the pLVAD group required a significantly lower rate of epinephrine at 0.13 (0.04) vs. 0.36 (0.06) mcg/kg/min (*p*=0.004) and norepinephrine at 0.32 (0.05) vs. 0.56 (0.07) mcg/kg/min (*p*=0.014) at 30 minutes post ROSC when compared to the Standard care group to maintain a MAP of greater than 70 mmHg and cardiac output of greater than 2.0 L/min (**Figure 6**).

**Figure 6.**
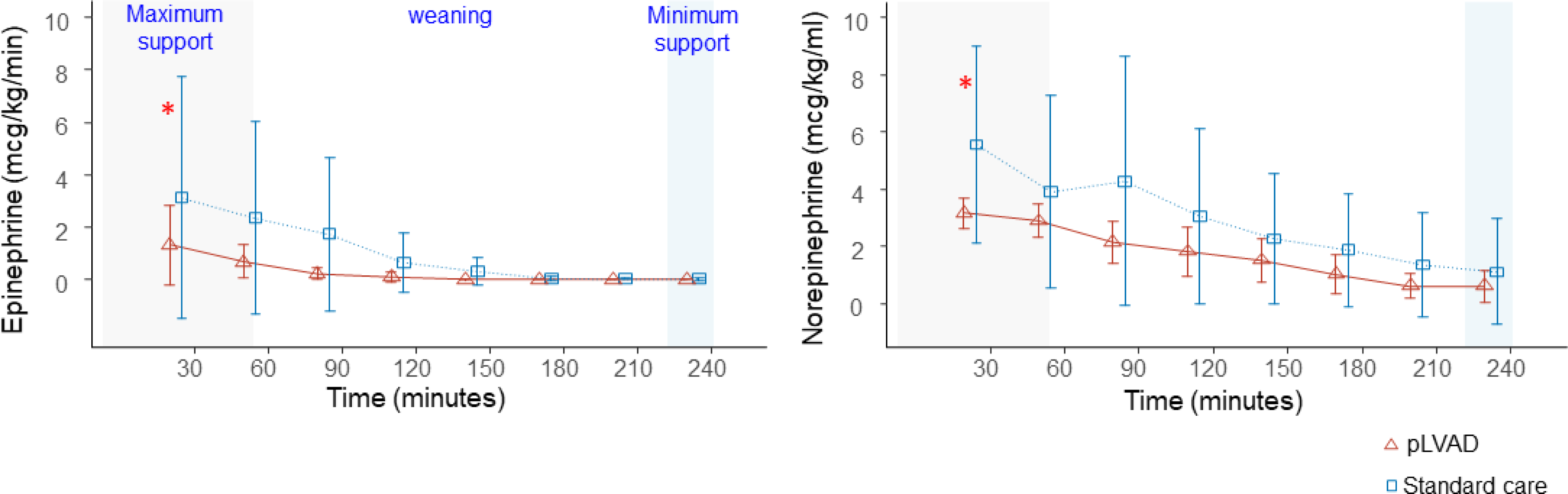
Longitudinal analyses of vasopressor after resuscitation. Red and blue lines indicate pLVAD group and Standard care group, respectively. * statistically significance (p <0.05) ** statistically significance (p<0.01) pLVAD denotes percutaneous left ventricular assist device.

## Discussion

Our study demonstrated that, among swine models with prolonged cardiac arrest, animals treated with pLVAD had higher CI and LVEF when compared with swine treated with standard of care at 240 minutes after ROSC. This is the first study showing the effect of pLVAD support on the recovery of cardiac function after prolonged cardiac arrest.

Refractory shock due to post-cardiac arrest myocardial dysfunction is a major cause of cardiac arrest recurrence within the first 24 hours after ROSC.^7, 10, 19, 20^ In one series of consecutive 148 patients admitted after successful ROSC after OHCA, 49% of patients had hemodynamic instability with median LVEF of 32%.^7^ In another study, which includes 47 VF-OHCA patients with therapeutic hypothermia and pulmonary artery catheter monitoring, 66% of the patients had post-arrest myocardial dysfunction with a median CI of 1.25 L/min/m^2^.^10^ In the present study, the standard care group had hemodynamic instability with mean EF of 32% and mean CI of 2.3 L/min/m^2^ at 30 minutes after ROSC, respectively, despite high dose of vasopressor. Our prolonged cardiac arrest model was similar in post-cardiac arrest cardiac dysfunction to the clinical scenarios.

ECPR is effective as both a rescue therapy for patients who fail to achieve ROSC but also as a hemodynamic stabilization therapy for refractory shock after ROSC. However, the effect of veno-arterial extracorporeal membrane oxygenation (V-A ECMO) on the recovery of myocardial function is controversial based on the risk of increased LV afterload which could exacerbate post-arrest myocardial dysfunction.^21–24^ Our recent study using a swine model of prolonged cardiac arrest treated with ECPR demonstrated that only 65% (15 /23) of subjects that achieved ROSC were successfully weaned from ECMO, defined as less than 1 L/min support. At 480 minutes after ROSC, the average LVEF was 50 (4) %, which corresponded to 77 (6) % of baseline.^25^ Given that all animals with ROSC in the present study were weaned from pLVAD at 240 minutes with 99 (11) % recovery of LVEF, future research directly comparing post-cardiac arrest VA-ECMO support and pLVAD support is warranted.

Focusing on the early post-ROSC period during which maximum pLVAD flow was maintained provides a potential mechanism by which pLVAD support was associated with better recovery of myocardial function. Initially, pLVAD significantly reduced the dose of inotrope /vasopressor by providing hemodynamic stability. High dose of these agents have been shown to increase vascular resistance and myocardial oxygen consumption.^26^ Animal^27, 28^ and clinical studies^7^ have demonstrated that high doses of epinephrine exacerbated post-arrest myocardial dysfunction. Subsequently, pLVAD with high flow resulted in a significant elevation of CPP, despite no statistically significant difference in MAP between the groups. Previous animal studies, using an isolated canine heart model, demonstrated that a reduction of CPP deteriorates myocardial contractility and that this can be reversed through an elevation of CPP.^29, 30^ Additionally, pLVAD with high flow significantly reduced PCWP and LVSWI. Saku et al., using a dog model with acute myocardial infarction, reported that LV mechanical unloading by pLVAD in the acute phase significantly reduced infarction size.^11^ They posited that the reduction of myocardial oxygen consumption played a contributory role in the reduction of ischemic reperfusion injury. Our current findings are consistent with their results since there is a positive correlation between LVSWI and myocardial oxygen consumption.^31^ pLVAD also significantly reduced TPRI, while there were no statistically differences in PA compliance. The right ventricular (RV) is highly afterload sensitive.^32, 33^ Our results suggest that pLVAD decreased the afterload against RV under similar PA compliance, which may contribute to the earlier recovery of post-arrest RV dysfunction. However, since pLVAD was activated during CPR, we were not able to discern the extent to which intra-arrest LV unloading contributed to the recovery of post-arrest myocardial dysfunction.

### Study Limitations

Our study has several limitations. First, pLVAD was initiated during CPR and the benefit of pLAD during CPR may have also contributed to recovery of myocardial function. Second, myocardial oxygen consumption, coronary perfusion flow and LV/RV pressures could not be measured directly. Finally, we did not measure long term outcomes, and therefore the cardiac function of animals treated with standard care might be delayed and gradually recover in the long term.

## Conclusion

Our animal study using a swine model of prolonged cardiac arrest demonstrated that early pLVAD support after ROSC was associated with better recovery of post-cardiac arrest myocardial function compared to standard of care. Further research is warranted to identify the indications and optimal strategies for post-ROSC mechanical circulatory support.

